# A thermodynamic analysis of CLC transporter dimerization in lipid bilayers

**DOI:** 10.1101/2023.03.14.532678

**Authors:** Rahul Chadda, Taeho Lee, Priyanka Sandal, Robyn Mahoney-Kruszka, Janice L. Robertson

**Affiliations:** Department of Biochemistry and Molecular Biophysics, Washington University School of Medicine, St. Louis, MO; Department of Physics, Washington University, St. Louis, MO; Department of Molecular Physiology and Biophysics, The University of Iowa, Iowa City, IA

## Abstract

The CLC-ec1 chloride/proton antiporter is a membrane embedded homodimer where subunits can dissociate and associate, but the thermodynamic driving forces favor the assembled form at biological densities. Yet, the physical reasons for this stability are confounding since binding occurs via the burial of hydrophobic protein interfaces yet the hydrophobic effect should not apply since there is little water within the membrane. To investigate this further, we quantified the thermodynamic changes associated with CLC dimerization in membranes by carrying out a van ′t Hoff analysis of the temperature dependency of the free energy of dimerization, *ΔG°*. To ensure that the reaction reached equilibrium under changing conditions, we utilized a Förster Resonance Energy Transfer based assay to report on the relaxation kinetics of subunit exchange as a function of temperature. These equilibration times were then applied to measure CLC-ec1 dimerization isotherms as a function of temperature using the single-molecule subunit-capture photobleaching analysis approach. The results demonstrate that the dimerization free energy of CLC in *E. coli* membranes exhibits a non-linear temperature dependency corresponding to a large, negative change in heat capacity, a signature of solvent ordering effects including the hydrophobic effect. Consolidating this with our previous molecular analyses suggests that the non-bilayer defect required to solvate the monomeric state is the molecular source of this large change in heat capacity and is a major and generalizable driving force for protein association in membranes.

## INTRODUCTION

A long-standing mystery surrounding membrane proteins is how they fold and form stable macromolecular complexes inside the hydrophobic environment of the cellular membrane. As described in the two-stage model of alpha-helical membrane protein folding (1) non-polar helices must first partition into the membrane phase, and then these helices come together to form a thermodynamically stable folded structure, often via non-polar interfaces. The sheer observation of these structures indicates that the folded/assembled ensemble must be thermodynamically favorable over dissociated and disassembled forms, where the non-polar interfaces are solvated by the surrounding non-polar lipid solvent. For soluble proteins, which also assemble via hydrophobic interfaces, association is driven in large part due to the gain in free energy upon the burial of non-polar residues away from water, a phenomenon known as the hydrophobic effect (2). Yet, in the membrane, there is very little water, and so we do not expect the hydrophobic effect to be relevant. Why then is the free energy of the assembled state more favorable in membrane environment and is there a compensatory generalizable solvent-dependent driving force that takes the place of the normally ubiquitous hydrophobic effect? This remains one of the major unanswered questions in our understanding of protein folding due to the limited amount of thermodynamic information of membrane protein assembly reactions in membranes.

To address this gap in our knowledge, we established a robust model system where greasy membrane protein association equilibrium is quantifiable in lipid bilayers (3–8). CLC-ec1 is a Cl^-^/H^+^ antiporter that forms a homodimer of two identical subunits, each containing its own independent transport pathway (9). The non-polar dimerization interface is relatively large, 1200 Å^2^ per subunit and embedded within the membrane’s hydrocarbon core. It is a robust protein system where the integrity of functional fold can be measured using reconstituted chloride transport assays (10), and structure can be determined using x-ray crystallography (11). It has also been demonstrated to be capable of existing in both dimeric and monomeric forms while preserving structure and function (3). Following this, we developed methods for quantifying the in-membrane equilibrium dimerization reaction of the CLC antiporter using single-molecule microscopy (4). This approach demonstrates that the dimerization is reversible and yields a free energy of stability of -10.9 kcal/mole relative to the 1 subunit/lipid standard state, in 2:1 POPE/POPG lipid bilayers. To put this into a practical perspective, this means that 10 subunits expressed in the inner membrane of *E. coli* containing 10^7^ lipids, will be observed in the dimeric form 90% of the time. From a biological standpoint, we interpret this thermodynamic stability as being more than sufficient as the dimer will be the predominant form at any reasonable expression level. Thus, it is apparent that CLC has evolved a mechanism for strong stability within the membrane phase via greasy interfaces alone, while the physical reasons for this stability remain unknown.

A critical step in identifying the physical and molecular driving forces involved is the ability to decompose free energies into thermodynamically meaningful changes in enthalpy and entropy. In general, changes in enthalpy reflect differentials in non-bonded interactions, while changes in entropy reflect the difference in accessible microstates for the system. Understanding how each of these thermodynamic parameters change, in tandem with strategic inspection of the involvement of the protein and solvent, can reveal the molecular factors that define the balance of equilibrium. One way of accessing this information is by carrying out a van ′t Hoff analysis of the temperature dependency of the reaction free energy. The change in the standard-state Gibbs free energy associated with dimerization, *ΔG°*, is defined as:

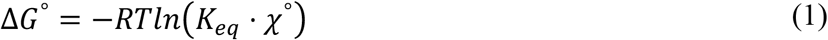

where *R* is the gas constant, *T* is the temperature in Kelvin, *K*_*eq*_ is the equilibrium association constant and *χ*° is the standard-state, which we define in mole fraction units as 1 subunit/lipid. Typically, the standard-state factor is omitted from this equation as it is conventionally set to the mathematically convenient value of 1 mole/L. However, since we are working in the 2-dimensional membrane reaction space (12) we select the similarly arbitrary, but mathematically convenient standard-state of 1 subunit/lipid for in-membrane reactions. Note, the term is explicitly included in eq. 1 so it is not forgotten, especially since other standard state definitions, such as area density are often used in the literature in this field. At a given temperature, *T*, the free energy is also equal to:

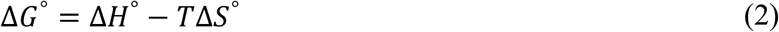

where *ΔH°* and *ΔS°* are the changes in standard-state enthalpy, and entropy associated with the dimerization reaction. Equating (2) and (3), yields the van ′t Hoff equation:

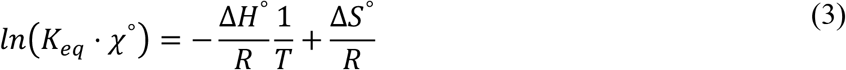

and plotting *ln* (*K*_*eq*_ · *χ* °) vs. *1/T* allows for the determination of *ΔH°* and *ΔS°*. If the enthalpy and entropy changes are temperature independent, then the relationship will be linear, with slope = -*ΔH°/R* and y-intercept = *ΔS°/R*. However, proteins and their surrounding solvent, e.g., water or lipids, can possess high heat capacities, meaning that there may be significant changes in heat capacity associated with changes in protein conformation or assembly. With this, enthalpy and entropy changes will also depend on the temperature as follows:

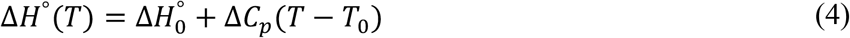

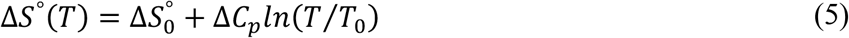

where *ΔC*_*P*_ is the change in molar heat capacity, *T*_*0*_ is an arbitrary reference temperature, and *ΔH*_*0*_*°* and *ΔS*_*0*_*°* are the respective changes in enthalpy and entropy at the reference temperature. Thus, combining eq. (4) and (5) with (3), the van ′t Hoff equation becomes a non-linear relationship:

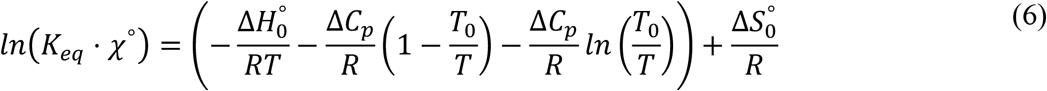

This is a complex function that has a quadratic dependency, but non-linear curve-fitting of *ln* (*K*_*eq*_ · *χ* °) vs. *1/T* with eq. 6 can provide the thermodynamic changes *ΔC*_*P*_, *ΔH*_*0*_*°, ΔS*_*0*_*°* and *T*_*0*_ that accompany the reaction.

In the present study, we report a thermodynamic dissection of the CLC-ec1 dimerization equilibrium in *E. coli* polar lipid (EPL) bilayers by carrying out a van ′t Hoff analysis of the temperature dependency of the dimerization free energy. The reason for changing to EPL is because the membranes are expected to be in liquid disordered phase over a wider range of temperatures, based on reported phase transition temperatures: *T*_*m,EPL*_ ≈ 2 °C (13) vs. *T*_*m,2:1POPE/POPG*_ ≈ 19 °C (14). To ensure equilibration of our samples across a wide range of temperatures, we developed a dynamic in-membrane subunit-exchange kinetic assay by measuring time-dependent ensemble FRET. In addition, we verify that the protein remains functionally capable of chloride transport from 22 - 62 °C under the same incubation conditions. With this, we can measure equilibrium CLC-ec1 dimerization isotherms as a function of temperature using the single-molecule subunit-capture photobleaching approach. The results show that the free energy of CLC-ec1 dimerization in EPL membranes is temperature dependent, with a non-linear van ′t Hoff relationship indicating a large negative change in heat capacity, *ΔC*_*P*_ = -2.5 kcal mol^-1^ K^-1^ and consequently, temperature dependent relationships of *ΔH°* and *ΔS°*. Such thermodynamic changes are a signature of the hydrophobic effect (15) or other solvent ordering such as those observed in phase transitions. Consolidating these results with our previous computational analysis of membrane solvation structures around the dimerization interface indicates that the non-bilayer membrane defect appearing exclusively in the monomeric state is the molecular basis for high heat capacity and the energetic driving force for dimerization in membranes.

## RESULTS

### CLC-ec1 dimers exhibit dynamic subunit exchange in lipid bilayers

Before measuring the temperature dependency of the dimerization equilibrium of CLC-ec1, we first needed a way of monitoring reaction equilibration as a function of temperature. Previously, we assessed the dynamic equilibration of CLC-ec1 dimerization in 2:1 POPE/POPG lipid bilayers using the single-molecule subunit capture approach (4). This method examines the population of Cy5 labeled CLC-ec1 subunits that are reconstituted into large, multilamellar membranes. The samples can be incubated under different conditions, and then the protein population is assessed by measuring the random capture of subunits into liposomes, which follows Poisson-like statistics. This is quantified by direct counting of subunit occupancy in liposomes using single-molecule photobleaching analysis. Provided the protein density, Cy5 labeling yield and liposome size distribution are known, then the photobleaching probability distribution directly reports on the proportion of monomers and dimers in the original membranes. In our previous study, we examined time-dependent equilibration by carrying out a perturbative dilution study in the membrane, where a dimeric population was diluted by freeze-thaw fusion with empty vesicles and the protein population observed to shift towards monomers when examined by single-molecule photobleaching (4). This reaction was studied at room temperature, ≈ 22 °C, and the conversion of the photobleaching distribution, with increasing single-step photobleaching probabilities, *P*_*1*_, and decreasing two-step probabilities, *P*_*2*_, were observed to slowly relax with *t*_*1/2*_ = 4.2 days.

This study demonstrated path-independence of the reaction, but since it only reflects the shift towards monomers, we wanted to develop another method to report on the formation of new dimeric species in the system. We did this by measuring the subunit-exchange behavior of CLC-ec1 at constant protein density. In this assay, WT CLC-ec1 is labelled by either a FRET donor, Cy3, or FRET acceptor, Cy5, and then reconstituted in EPL bilayers at densities where the dimeric form prevails, e.g., 1 μg/mg or *χ* = 10^−5^ subunits/lipid (**Fig. 1A**). The donor and acceptor-labelled subunits are then introduced into the same membrane by freeze/thaw mediated fusion, enabling the dynamic exchange of subunits leading to the formation of new Cy3/Cy5 heterodimers was monitored by measuring the ratiometric FRET signal. As a negative control, the disulfide cross-linked form of CLC-ec1 was used, R230C/L249C (RCLC), that is incapable of subunit exchange (6, 16). Separately labelled proteoliposome samples of WT_Cy3_ and WT_Cy5_ were aliquoted in the same cuvette at a ratio of *P*_*Cy5*_*/P*_*Cy3*_ = 6.0 ± 0.5, but not fused, yielding a baseline FRET signal for WT_Cy3+Cy5_ of 0.20 ± 0.01. This reflects the background FRET signal that corresponds to zero Cy3/Cy5 heterodimers **(Fig. 1B, C)**. However, a positive control where WT is co-labelled with Cy3/Cy5 at a ratio of *P*_*Cy5*_*/P*_*Cy3*_ = 6.2 ± 0.6, yielded a FRET signal for WT_Cy3/Cy5_ of 0.32 ± 0.01, a significant increase over the background. Freeze-thaw fusion of the baseline samples into multi-lamellar vesicles (MLVs) introduces WT_Cy3_ and WT_Cy5_ dimers into the same bilayer, i.e., WT_Cy3+Cy5_^*fused*^. Upon incubation of WT_Cy3+Cy5_^*fused*^ sample at room temperature, a slow but consistent increase in FRET is observed which converges to the WT_Cy3/Cy5_ co-labelled FRET signal with a *t*_*1/2*_ = 8.9 ± 1.8 days **(Fig. 1D, E)**, on the order of what we observed in our photobleaching dilution study in 2:1 POPE/POPG (4). On the other hand, the constitutively dimeric RCLC shows unchanged FRET signals over the same period for both RCLC_Cy3+Cy5_^*fused*^ and co-labelled RCLC_Cy3/Cy5_ samples (5, 13) ruling out artifacts from non-specific protein aggregation or differential bleaching of fluorophores along this timescale. Altogether, these results support the hypothesis that the exponentially saturating increase in FRET signal observed in WT samples is due to CLC subunit-exchange within the membrane.

**Figure 1.**
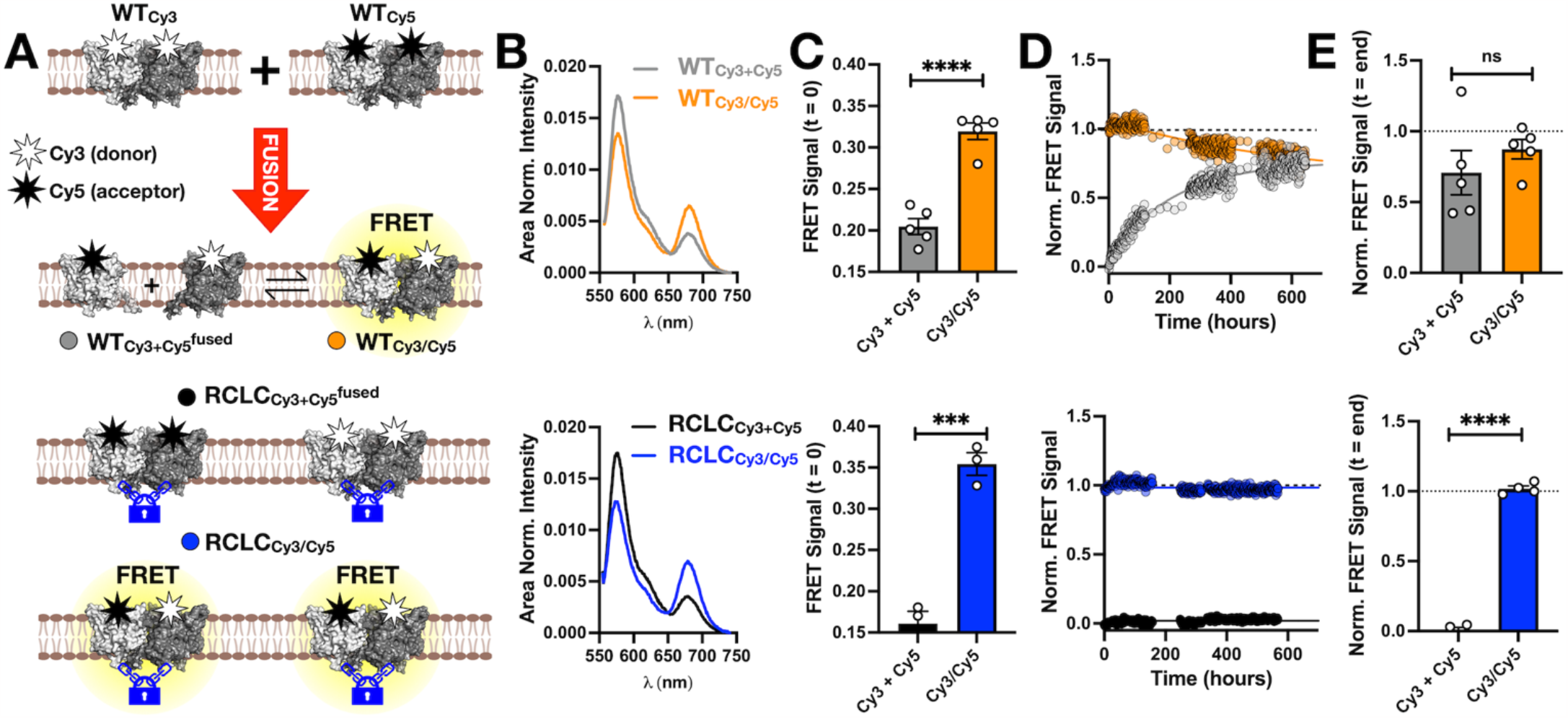
Dynamic exchange CLC-ec1 subunits in membranes observed by bulk FRET. **(A)** Cartoon depicting the subunit exchange reaction in membranes. CLC-ec1 is labelled with FRET donor Cy3 (white star) or FRET acceptor Cy5 (black star) and reconstituted into EPL bilayers at *χ* = 10^−5^ subunits/lipid. The following samples are studied: WT_Cy3+Cy5_ (light grey) indicates separately labelled proteoliposome samples that have been fused into the same membrane by 3x freeze-thaw, the co-labelled WT_Cy3/Cy5_ protein (orange), RCLC_Cy3/Cy5_ – disulfide cross-linked dimer CLC-ec1 R230C/L249C co-labelled with Cy3 and Cy5, and RCLC_Cy3+Cy5_ (black) disulfide cross-linked CLC-ec1-Cy3 and CLC-ec1-Cy5 incapable of subunit exchange, fused into the same membrane. **(B)** Area normalized bulk emission spectra from excitation at 565 nm for WT_Cy3+Cy5_, WT_Cy3/Cy5_, RCLC_Cy3+Cy5_ and RCLC_Cy3/Cy5_ right before fusion, defined as *t* = 0. **(C)** Initial FRET signal = *I*_665nm_/(*I*_565nm_ + *I*_665nm_) showing significant heterodimeric signals. Data represented as mean ± sem, n = 5 for WT, n = 3 for RCLC. Labeling yields of all constructs were comparable, *P*_*Cy5*_*/P*_*Cy3*_ = 3.1-7.6. Statistics were calculated using the unpaired parametric student’s t-test: WT (****, *p* < 0.0001) and RCLC (***, *p* = 0.0007). **(D)** Time-dependent normalized FRET signals for WT and RCLC samples at ambient temperature, ≈ 22 ºC. The data is normalized by defining the initial time point of the Cy3+Cy5 samples as 0 and the co-labelled Cy3/Cy5 signal as 1. WT time-courses are fit to an exponential association function, while RCLC time-course are fit with a flat line. **(E)** Endpoint normalized FRET signals, defined from the plateau value for the exponential fit, for WT (ns, *p* = 0.36) and RCLC (****, *p* < 0.0001). Data represented as mean ± sem, n = 5 for WT, n = 4 for RCLC.

### CLC-ec1 undergoes subunit-exchange across many temperatures while remaining functionally folded

With evidence of dynamic subunit exchange of CLC-ec1 in membranes at room temperature, we carried out the same experiment while incubating samples at different temperatures from 31 to 80 °C. As expected, increasing temperature accelerates the rate of CLC subunit exchange. From 31 - 44 °C, we observe comparable behavior in the saturation of the FRET signals as we observed for the room temperature samples, with the WT_Cy3+Cy5_^*fused*^ signal converging to that of the WT_Cy3/Cy5_ co-labelled samples (**Fig. 2A, B**). However, at higher temperatures, the FRET traces no longer exhibit single component kinetics. At and above 44 °C, WT_Cy3/Cy5_ show biphasic behavior i.e., an initial, relatively rapid, decay in FRET followed by a plateau that converges to the endpoint WT_Cy3+Cy5_^*fused*^ signal (**see Fig. 2; panel 3**). However, at 62 °C and above, the second phase (or the plateau from 44 °C) also begins to show time dependency namely an increasing FRET signal in both WT and RCLC samples. For the WT samples, these two behaviors can be discerned by fitting with fast and slow time dependencies (**Fig. 2B**), whereas the RCLC samples only show the slow time component. Examining the plateau FRET values from these fits shows that the slow component leads to an increase in FRET beyond the co-labelled WT_Cy3/Cy5_ or RCLC_Cy3/Cy5_ dimeric samples suggesting possible higher-order aggregation of the protein within the membrane at temperatures > 62 °C. However, plotting the temperature dependency of the fast time component between 22 - 62 °C shows a linear relationship between the logarithm of *t*_*1/2, fast*_ and temperature (**Fig. 2D**), in line with an Arrhenius model describing an equilibrium reaction in membranes. As a final test, we examined whether the increasing FRET signal observed in WT_Cy3+Cy5_^*fused*^ samples at 37 °C was reversible by diluting the end-point FRET samples by freeze-thaw fusion with 5-times excess of un-labelled WT proteoliposomes. Upon fusion, we observe that both WT_Cy3+Cy5_ and WT_Cy3/Cy5_ signals exponentially decrease (**Fig. 2E, F**) while RCLC samples show no change (**Fig. 2G, H**). Altogether, these results support the hypothesis that the fast time dependency in FRET signals observed from 22 - 62 °C reflects the dynamics of CLC subunit exchange, and that this behavior is separable from other reactions, such as aggregation, that dominates at higher temperatures.

**Figure 2.**
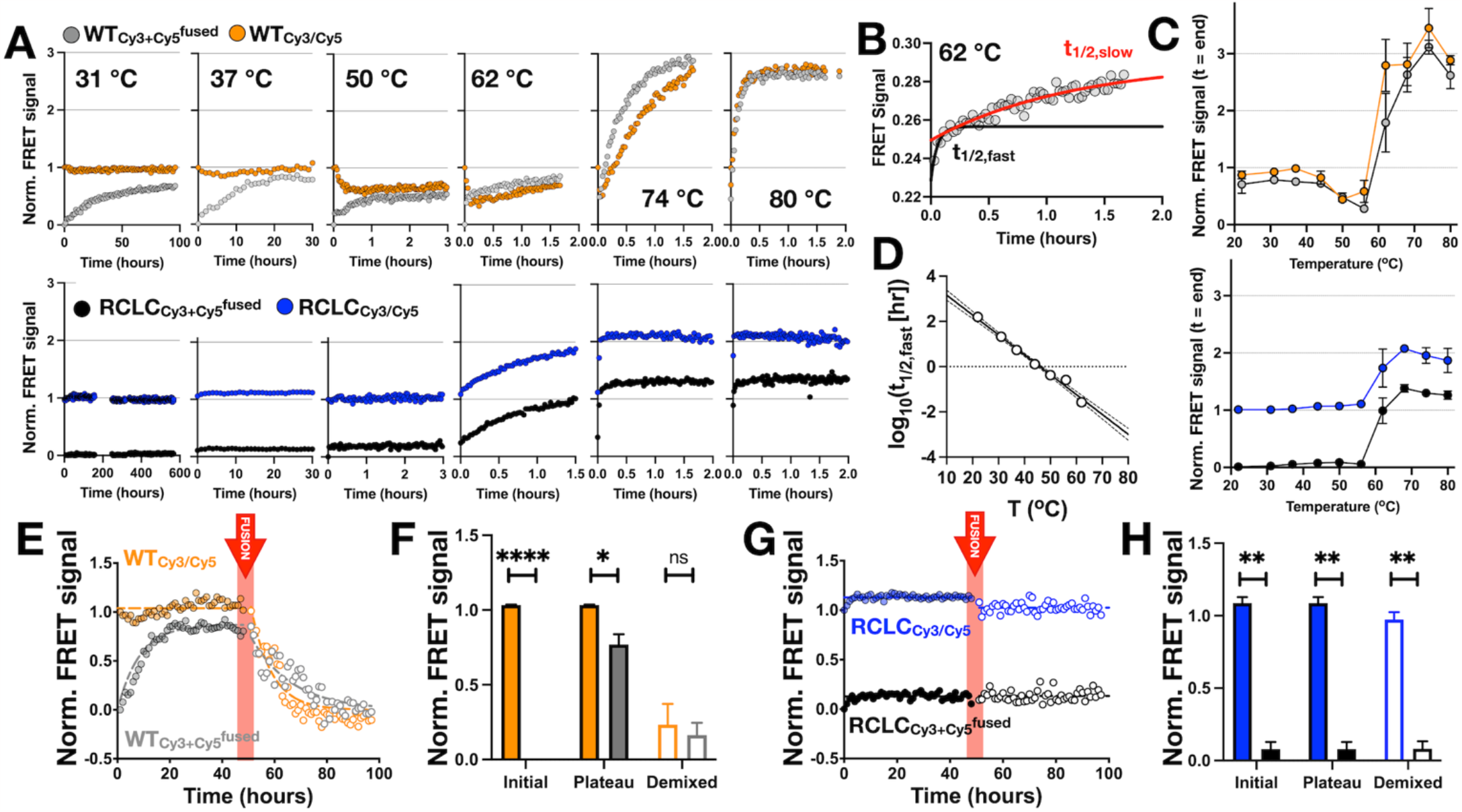
Temperature dependency and reversibility of CLC-ec1 subunit exchange in membranes. **(A)** Normalized time-dependent FRET signals for WT and RCLC samples as a function of increasing incubation temperature, 31 ºC – 80 ºC. **(B)** Fitting of time-dependent FRET signals of WT _Cy3+Cy5_ ^*fused*^ with a two-phase exponential association function, *t*_*1/2, fast*_ (red) and *t*_*1/2, slow*_ (green). **(C)** Normalized end-point FRET signals from the fitted plateaus. Data represented as mean ± sem, n = 2-6. **(D)** Logarithmic plot of *t*_*1/2, fast*_ as a function of temperature. **(E)** Reversibility of the heterodimeric WT_Cy3/Cy5_ FRET signal at 37 °C examined by freeze/thaw fusion with 5x volume of unlabeled WT proteoliposomes. **(F)** End-point FRET signals before and after dilution by unlabeled protein. **(G)** Disulfide cross-linked RCLC_Cy3/Cy5_ does not show reversibility, **(H)** with no significant changes in end-point FRET signals after dilution.

As a final test, we investigated whether the subunit-exchange exists between folded subunits by measuring chloride transport function from the exact same proteoliposomes studied in our FRET measurements, after the equilibration was complete, about 8 x *t*_*1/2, fast*_ (**Fig. 3A**). After incubation at elevated temperatures, the membranes are still capable of holding a chloride gradient (**Fig. 3B**) and maintain the capacity to transport chloride at comparable rates (**Fig. 3C**) with similar fractions of active vesicles (**Fig. 3D**) compared to liposomes at room temperature. Only samples that were incubated > 62 °C and for extended periods of time showed a significant decrease in activity corresponding to possibly aggregated protein accompanied by the increased FRET signals (**Fig. 2A, C**). Similar results were observed with the RCLC controls (**Fig. 3 – Supp. 1**). Finally, we measured the transport activity of WT directly at 37 °C, and the transport activity increases, further supporting the model that elevated temperatures accelerate the kinetics of exchange of functionally folded subunits in a dimeric assembly.

**Figure 3.**
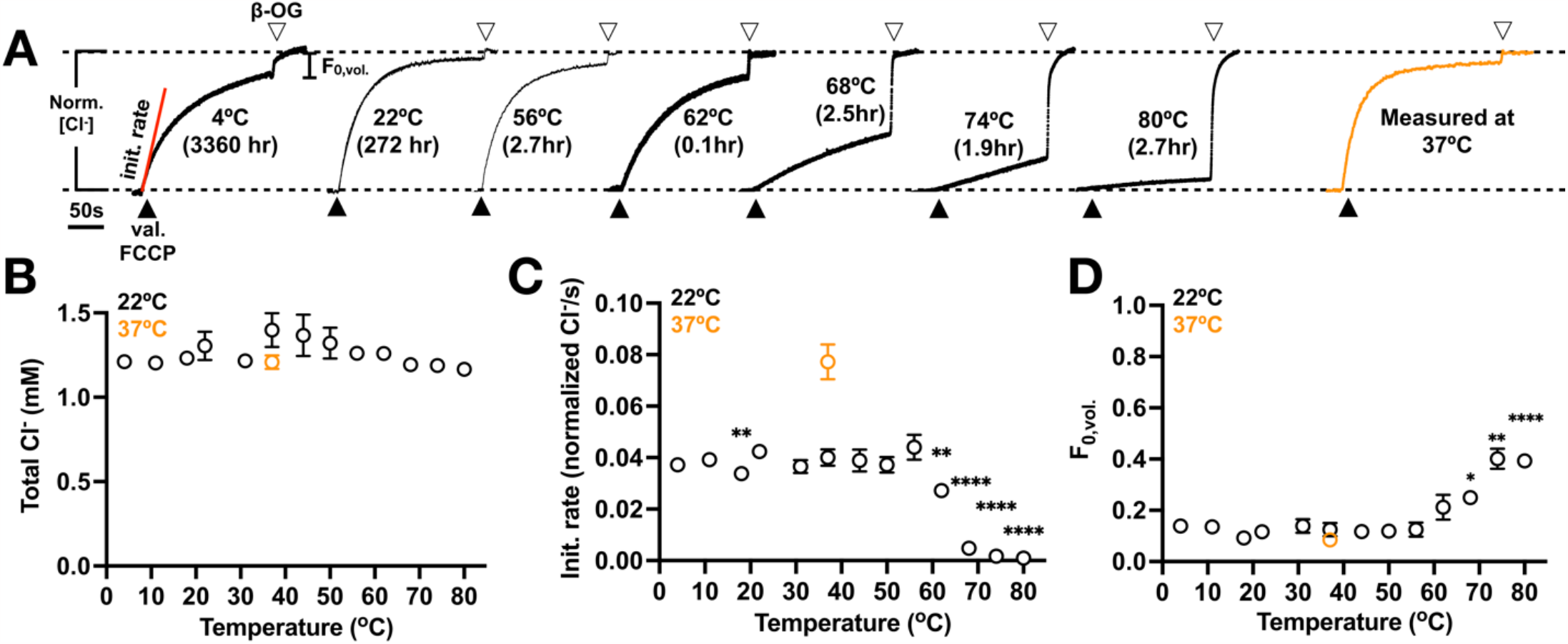
CLC-ec1 remains functional after incubation up to 62 ºC. **(A)** Representative chloride efflux traces from liposomes reconstituted at *χ* = 4 × 10^−6^ subunits/lipid in EPL incubated at 4-80 ºC for the indicated durations and then measured at room temperature. Black triangle indicates the addition of valinomycin and FCCP, white triangle indicates the addition of β-OG. The orange trace indicates a chloride transport measurement conducted directly at 37 ºC. **(B)** Total Cl^-^ concentration in the measurement cell after addition of β-OG, **(C)** initial chloride efflux rate, and **(D)** fractional volume of inactive vesicles, *F*_*0, vol*._ as a function of incubation temperature. Data represent mean ± sem, *n* = 2-4 and statistical tests were calculated out using an un-paired parametric student’s t-test compared to the 22 ºC data. The incubation time of the samples are 3360 ± 0 hr (4 ºC, 11 ºC, 18 ºC), 272 ± 128 hr (22 ºC), 71.7 ± 13.9 hr (31 ºC), 35.3 ± 7.4 hr (37 ºC), 16.6 ± 6.7 hr (44 ºC), 6.9 ± 1.3 hr (50 ºC), 2.7 ± 0.2 hr (56 ºC), 0.1 ± 0 hr (62 ºC), 2.5 ± 0.4 hr (68 ºC), 1.9 ± 0.1 hr (74 ºC) and 2.7 ± 0.7 hr (80 ºC) (mean ± sem, n = 2-4) and chloride transports are measured at 22 ºC.

### The temperature dependency of CLC-ec1 dimerization equilibrium in EPL membranes

Having validated that the CLC dimerization reaction is well-defined from 22 - 62 °C, we carried out a van ′t Hoff analysis of the dimerization free energy by measuring equilibrium dimerization isotherms across the pre-validated range of incubation temperatures. From our previous experiments monitoring time dependent FRET, we observed some changes in the plateau FRET values that could correspond to changes in dimerization. To quantify the stability changes rigorously, we returned to our single-molecule subunit capture approach that allows us to carry out a full titration of the dimerization reaction as a function of protein density to measure the equilibrium constant as a function of temperature (4, 5). In these studies, we reconstituted the protein across a density range of *χ* = 10^−9^ to 10^−6^ subunits/lipid, again in the membrane condition of *E. coli* polar lipid (EPL) extract. This composition contains approximately 67% PE, 23% PG, 10% cardiolipin, but has a lower phase transition temperature, *T*_*m*_, compared to pure 2:1 POPE/POPG (13, 14), ensuring that the membrane is in the liquid disordered state across a wider temperature range. We find that the EPL lipid environment has a stabilizing effect on the CLC-ec1 dimerization, with *K*_*eq*_ = 3.5 ± 2.6 × 10^9^ lipids/subunit and *ΔG°*_*EPL*_ = -12.7 kcal/mole, resulting in a stabilization of *ΔΔG*_*EPL-2:1 POPE/POPG*_ = -1.8 kcal/mole.

To isolate and study the effect of temperature on the *K*_*eq*_, proteoliposomes prepared at *χ* = 10^−5^ subunits/lipid, i.e., 1 μg/mg density, were serially diluted by freeze/thaw fusion with excess membranes and then incubated at various temperatures for the appropriate equilibration time (**Fig. 4A**). This perturbative approach allowed ensured identical initial conditions for all samples, and with that an ability to robustly measure changes in the dimerization isotherms due to the difference incubation conditions. The sample diluted and incubated at 22 °C showed no change in photobleaching probability distribution over period of 1-3 days acting as a non-equilibrated negative control (**Fig. 4B**). For the other temperatures, 31 - 62 °C, the samples are incubated for ≈ 8 x *t*_*1/2, fast*_ as measured by the bulk FRET subunit-exchange relaxation kinetics. The results show that the equilibrium dimer distribution shifts with temperature, which is apparent above 53 °C, reflected in the distribution of the dimerization free energies (**Fig. 4C**). Plotting the data in van ′t Hoff form, *ln* (*K*_*eq*_ · *χ* °) vs. *1/T*, shows that it is best supported by a non-linear van ′t Hoff model, indicative of a change of heat capacity upon dimerization (**Fig. 4D**). Over the temperature range of *T* = 22 - 62 *°C*, this yields a reference temperature of *T*_*0*_ = 300.0 ± 1.6 K and thermodynamic parameters of *ΔH*_*0*_*°* = 10.0 ± 4.1 kcal mol^-1^ K^-1^, *ΔS*_*0*_*°* = 0.077 ± 0.014 kcal mol^-1^ K^-1^, and *ΔC*_*P*_ = -2.5 ± 0.2 kcal mol^-1^ K^-1^ (**Fig. 4E**). The large change in heat capacity consequently means that *ΔH° and ΔS°* are temperature dependent functions (**Fig. 4F**), and at the physiological temperature of *T* = 37 *°C, ΔH°*_*310*_ = -15 kcal mol^-1^ and *-TΔS°*_*310*_ = 1.5 kcal mol^-1^. A consequence of a non-linear van ′t Hoff relationship is that it exhibits a quadratic appearing dependency on temperature, predicting that dimerization becomes weaker at colder temperatures. To test this, we attempted measurements below room temperature, but did not observe significant perturbations in our starting probability distributions, which we interpret is due to the exceedingly slow kinetics under these under these conditions (**Fig. 4 – supplement 1**). Therefore, we limit our conclusions for this reaction to the testable range of *T* = 22 – 62 *°C*.

**Figure 4.**
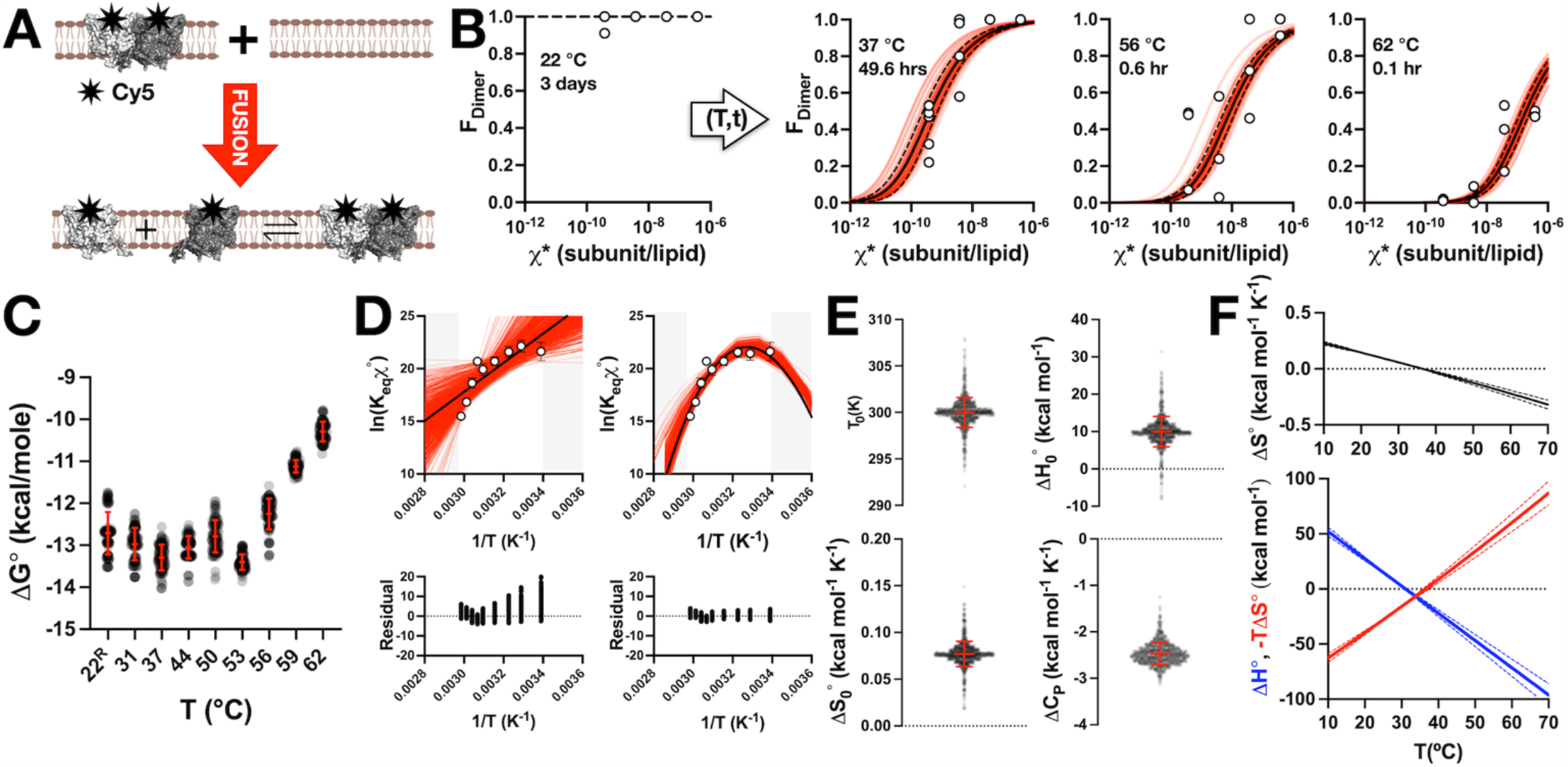
The temperature dependency of CLC-ec1 dimerization equilibrium in EPL membranes fits a non-linear model indicative of a large change in heat capacity upon association. **(A)** Schematic of the dilution titration experiment. WT_Cy5_ CLC-ec1 in reconstituted in EPL at density of *χ*_rec._ = 2 × 10^−5^ subunits/lipid where dimers are predominant. The final mole fraction density is achieved by dilution in the membrane 10-10,000x by freeze-thaw fusion with empty EPL vesicles, then incubated at the indicated temperature for t_inc._ ≈ 8 x t_1/2_ of the subunit-exchange relaxation time and then examined using the single-molecule photobleaching subunit capture approach. **(B)** Resultant dimer population, *F*_*Dimer*_ vs. *χ*\*, for non-equilibrated samples at 22 ºC diluted an assessed after only 3 days incubation, 37ºC diluted after 49.6 hr incubation, 56 ºC dilution after 0.6 hr, and 62 ºC dilution after 0.1 hr incubation. Data represent *n* = 3-5 independently prepared samples. Red lines indicate bootstrap fitting of the experimental data to the equilibrium dimerization isotherm in MATLAB, with 8 data points selected at random from a total set of 12-20, iterated 100 times and limits = {10^−10^, 10^−5^}. Black line indicates direct non-linear fitting of all experimental data to the dimerization isotherm with GraphPad Prism. **(C)** Bootstrapped distributions of *ΔG°* (kcal/mole). All populations are significantly different (*p* < 0.0001, un-paired parametric t-test) from the reconstituted, equilibrated 22 ºC sample (*22*^*R*^) except for the 50 ºC sample (*p* = 0.23). **(D)** Linear (*In K*_*eq*_ (T) = − Δ*H* ° / *RT* + Δ*S*° / *R*) vs. non-linear 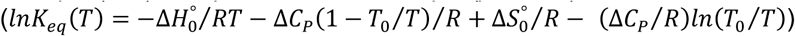 van ′t Hoff fitting results carried out by bootstrapping analysis in MATLAB with 15 data points selected from a total set of 900, iterated 1000 times. Residuals are plotted below each graph. **(E)** Distributions of thermodynamic parameters from the non-linear bootstrapping fits (mean ± standard deviation over 1000 bootstrapped fits): *T*_*0*_ = 300.0 ± 1.6 K, *ΔH*_*0*_*°* = 9.98 ± 4.06 kcal mol^-1^, *ΔS*_*0*_*°* = 0.077 ± 0.014 kcal mol^-1^ K^-1^ and *ΔC*_*P*_ = -2.47 ± 0.24 kcal mol^-1^ K^-1^. **(F)** Temperature dependent enthalpy, 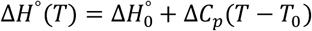, blue, and entropy, 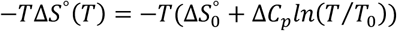, red, contributions to the change in free energy of CLC dimerization in membranes. Data represented as mean ± standard deviation over 1000 bootstrapped values.

## DISCUSSION

### CLC dimerization is in a dynamic equilibrium in lipid membranes

Adding to our previous studies (4–8), the results presented here demonstrate that the large CLC-ec1 chloride/proton antiporter participates in a dynamic dimerization reaction inside of the lipid bilayer. This involves dissociation and re-association of the subunits inside the membrane all while maintaining functional transport activity. At the physiological condition of 37 *°C*, the heterodimeric FRET time-course exhibits a half-time to steady state at about 6 hours, indicating that the reaction is slow but dynamic. Slow kinetics of membrane protein association within membranes has been observed before, as indicated in steric-trapping studies of diacylglycerol kinase trimers (17) and glycophorin-A dimers (18, 19). Since subunit-exchange kinetics are dependent on the reaction association and dissociation rates, and other variables such as donor and acceptor stoichiometries (20), we reserve the quantitative analysis of the reaction kinetics for future study. Still, the observation of the exponentially increasing FRET signals and convergence to heterodimeric controls provides strong evidence that the FRET signal is reporting on the formation of heterodimers arising from dynamic subunit-exchange in the membrane, challenging the long-held presumption that CLC antiporters are obligatory complexes.

The evidence of dynamic assembly in membranes means that the probability of dimers is dependent on thermodynamically relevant parameters such as density of protein relative to lipid solvent, the composition of the lipid solvent, temperature, pressure, and chemical linkage. This means that the oligomerization is contextual and can redistribute in a regulatory manner inside of the membrane, even after dimers have formed. We observe this through lipid dependencies on the dimerization equilibrium, for example, as is observed when we switch the membrane composition to EPL from the synthetic mimic of 2:1 POPE/POPG, which stabilizes dimerization by about 2 kcal mol^-1^. We have yet to identify the reason for this stabilizing effect but speculate that it is a factor particular to the native extract, such as an unexplored lipid component. Yet, the observation that this reaction can be shifted by lipids, and in the opposite direction of the effect of short-chain lipids that we observed previously (7), further demonstrates we are working with equilibrium reactions in the membrane where the membrane plays a role in defining the thermodynamic balance.

While quantitative stabilities are an absolute first step, to understand the thermodynamic forces involved, we must push the analysis further and dissect the change in free energy into changes in enthalpy and entropy. In aqueous solutions, this sort of dissection can be straightforward, as the direct heats of association can be measured by isothermal calorimetry. However, this approach is not possible for equilibrium association measurements within the membrane phase, as one cannot simply add membrane to the system and have it spontaneously incorporate with the pre-existing lipid bilayer. With this, the study of thermodynamic dissection of membrane protein association in lipid bilayers has been limited to only a few studies, where a temperature dependency of the free energy can be measured using a van ′t Hoff analysis.

The first partial study of this sort was carried out for Gramicidin dimerization, a Na^+^-selective ion channel across the membrane that is formed by the association of two pentadecapeptides, one in each leaflet of the membrane (21). The probability of functional dimer formation is determined by the dynamic observation of electrical currents measured using single-channel bilayer electrophysiology. The temperature dependency of the dimerization was measured in dioleoyl-lecithin/n-decane at *T* = 10, 25 & 40 *°C*. However, these three points are not sufficient to confidently discriminate between different van ′t Hoff models required for a full thermodynamic analysis.

The second study that has been reported involves the dimerization of poly-alanine/leucine transmembrane helix measured by the temperature dependency of FRET (22–24). This is a weak affinity dimer complex, and so the studies could be carried out using bulk FRET measurements, and more recently expanded to single-molecule FRET microscopy where dynamic dimer formation is readily observable. As far as we know, these sets of studies reflect the historical record of quantitative, temperature dependency measurements for assembly of membrane embedded proteins in membranes. Furthermore, the thermodynamic dissection of the free energy of CLC dimerization in membranes reveals itself to be the first large biological transport complex of a multi-helical folded protein where thermodynamic information can now be obtained and provides an essential next step to understanding the physical behavior of other biological membrane protein systems.

It is important to note that while our results indicate that we can measure the temperature dependency of this complex whilst in the membrane, this may not be experimentally accessible for all proteins. Because of the slow kinetics at ambient temperatures, we used a trick of incubating the system at higher temperatures, then locked in the oligomeric distribution by bringing the samples to room temperature where we could measure full dimerization isotherms using our single-molecule subunit-capture approach. Still, because many membrane protein complexes have been observed to be slow to dissociate (17), it may be possible to extend this measurement to other membrane protein systems.

### Dimerization of CLC involves a large negative change in heat capacity

The van ′t Hoff analysis for the CLC dimerization free energy in EPL lipid bilayers exhibits a non-linear temperature dependency indicating that the conversion from monomers to dimers involves a large negative change in heat capacity. The concept of a heat capacity change can be a difficult to consider for those outside of the field of protein self-assembly, but particularly in the study of membrane proteins since we have not many opportunities to consider data like this so far. Thus, it is worth some extra discussion here. In their book on biological thermodynamics, Dill & Bromberg simply state that the heat capacity “*describes the storage of energy (or enthalpy) in bonds that break or weaken with increasing temperature*” (2). To elaborate on this, heat energy that is applied to the sample is distributed into both kinetic and potential energy (25). The increase in kinetic energy results in faster velocities and therefore an increase in temperature. However, the heat energy can also be distributed into potential energy of the vibrations of molecular bonds. As a result, these bonds can break, allowing for increased rotational freedom without increasing the velocity of the molecule. Thus, systems that contain substantial networks of non-bonded interactions can possess a high heat capacity. A classic example of a substance with high molar heat capacity is water since it participates in an extensive hydrogen bonding network. Addition of heat changes the hydrogen bond distribution, weakening bonds and decreasing the order of molecules relative to each other. Consider the change in state of water from ice to liquid. A rough estimate of the melting of a protein-sized 30 Å ice crystal like a snowflake, is predicted to exhibit ≈ 5 kcal mol^-1^ K^-1^ change in heat capacity, resulting from only the changes in the network of hydrogen bonds (25).

With this, we go back and consider our observation that the change in heat capacity of CLC dimerization reaction is -2.5 kcal mol^-1^ K^-1^. This is a stunningly large value that is on the order of the upper limit of heat capacity changes, comparable to the phase transition melting of a protein-sized ice crystal. What could be the molecular explanation for such a large heat capacity difference between the monomeric and dimeric states of CLC in membranes? For this we turn to our understanding of heat capacity changes in soluble protein self-assembly. Large changes in heat capacity have been observed to accompany significant changes in the exposure of non-polar groups to water, a phenomenon that is better known as the hydrophobic effect. The molecular explanation of this has been attributed to the unfolded or dissociated state, leading to exposure of non-polar surfaces to water. The surrounding first shell of waters cannot form hydrogen bonds with the non-polar groups, and therefore adopt highly organized solvation structures, often referred to as “*icebergs*” around these exposed surfaces. Addition of heat can be distributed into the remaining hydrogen bonds, weakening them to promote disordering of the water at the interface. The expected contribution from the hydrophobic effect can be predicted by measuring non-polar molecule partitioning into water. For example, benzene partitioning into water exhibits a heat capacity change of 0.27 cal mol^-1^ K^-1^ Å^-2^ (26), and this value can be multiplied by the expected change in non-polar surface exposure to predict the heat capacity change between the dissociated/unfolded and associated/folded states (27). In general, these predictions agree well with experimental measurements for both protein folding and oligomerization in water, reaching heat capacity changes of -1 kcal mol^-1^ K^-1^.

However, this value is already considerably lower than what we observe for CLC dimerization in the membrane, suggesting that the hydrophobic effect cannot be the only cause behind our finding. To add to this, if we apply the same hydrophobic effect calculation to CLC, which exposes about 2400 Å^2^ of its binding interface upon dissociation, we only expect *ΔC*_*P*_ = -0.7 kcal mol^-1^ K^-1^ in the direction of association. Considering that we know this interface remains membrane embedded during the entire reaction, since the protein remains functional for transmembrane chloride transport, the actual amount that is predicted to come from the hydrophobic effect must be much smaller. This therefore eliminates the hydrophobic effect as the only cause, and we must once again remind ourselves that there are other molecular reasons for large changes in heat capacity such as phase transitions in water, or perhaps the lipid solvent. In fact, anomalously large changes in heat capacity are often observed in the binding of highly charged species, such as proteins to DNA. One example is the case of the F Factor Relaxase to single-stranded DNA, which exhibits *ΔC*_*P*_ = -3.3 kcal mol^-1^ K^-1^ in the direction of binding (28), expected to be the result of both the hydrophobic effect but also base stacking. With this, we predict that the molecular contributions for the large heat capacity change observed in CLC dimerization involves substantial molecular ordering of all the different components in the system.

What sort of structural changes could account for such differences in molecular ordering? Our previous investigation into the changes in membrane structure around the monomeric and dimeric states of CLC provide some clues as to what may be conferring this anomalous heat capacity change (7). Extensive coarse-grained molecular dynamics simulations sampling lipids around the reaction endpoints, revealed that the membrane thins and twist to solvate the exposed dimerization interface in the monomeric state. This change in membrane shape arises from the favorable energetics of matching the membrane to the hydrophobic surface of the protein, however, it comes at a cost in terms of the total free energy of the system. But in the typical palmityl-oleoyl lipids that comprise EPL, these longer lipids cannot adequately pack in this region, resulting in significant lipid tilting, a reduction of local lipid packing and increased water penetration near the acyl chains. Naturally, the partitioning of water into the membrane is the hydrophobic effect, but only 4-5 waters are estimated to penetrate the acyl chains per monomer (*personal communication with N. Bernhardt*), insufficient to account for the full change in heat capacity that we measure. Thus, we consider that there are other components to this, such as changes in the hydrogen bonding of lipid headgroups and even the strength of non-bonded van der Waals interactions. Since many lipids are impacted around the dimerization interface, we anticipate that the sum of smaller effects over many lipid and water molecules will account for the large change in heat capacity that is observed in our system.

Finally, membrane dependent changes in heat capacity have been observed before, in the one other system where van ′t Hoff analysis was carried out on an in-membrane dimerization reaction (22, 23). In this study, the dimerization free energy of a synthetic transmembrane helix (AALALAA)_3_ was measured in C14 to C20 PC membranes exhibiting linear temperature dependencies. However, changing the lipid composition to the longer C22 lipids, which are hydrophobically mismatched to the dimer state, introduces a non-linear temperature dependency with a heat capacity change of +0.6 kcal mol^-1^ K^-1^ in the direction of dimerization. This implies that the monomeric state is more suitably solvated in the lipid bilayer, with the dimeric state ordering the solvent molecules in the system. The number of studies here are small, but both transmembrane helix dimerization and large CLC dimerization appear to have the capacity for changes in heat capacity linked to changes in solvent order factoring into the thermodynamic stability of dimerization within the membrane. Note the fact that both association reactions introduce non-linear temperature dependencies with large heat capacity changes suggests a potential mechanism for temperature sensitivity that may be pertinent to other membrane protein reactions within the membrane (29).

## CONCLUSION

This study demonstrates that CLC dimers are dynamic and reversible assemblies in the membrane, forming across a wide range of temperatures. This finding enables the rare thermodynamic dissection of the free energy for binding within the membrane by van ′t Hoff analysis. The results demonstrate that dimerization of the hydrophobic CLC interface is accompanied by a large negative change in heat capacity, resembling the signature of the hydrophobic effect in driving soluble protein dimerization in water. Placing these results in context with previous computational work suggests that this heat capacity change arises from the removal of a thinned membrane defect that is required to solvate the exposed dimerization interface in the monomeric state, which becomes buried upon binding. Finally, we find that the temperature dependency of CLC dimerization is pronounced, introducing a new model system for considering other temperature dependent reactions in the membrane such as ion channel gating.

## ACKNOWLEDGEMENTS

The Robertson lab is supported by the National Institute of General Medical Science, National Institutes of Health (R01GM120260, R21GM126476). We thank Tim Lohman, Nathan Bernhardt, and Chris Miller for useful discussions.

## MATERIALS AND METHODS

### Protein purification

DNA constructs for CLC-ec1 C85A/H234C (WT), and C85A/H234C/R230C/L249C (RCLC) have been described earlier, along with their expression and purification methods (4, 6). Briefly, BL21-AI *E. coli* cells transformed with the expression plasmid were lysed by sonication and the protein extracted from membrane fragments into 2% n-Decyl-β-D-Maltopyranoside (DM; Anatrace, Maumee OH) containing 5 mM TCEP (Tris (2-carboxyethyl) phosphine; Soltec Bioscience, Beverly, MA) to ensure that the introduced cysteine residue at residue 234 (H234C) remains in a reduced state for maleimide labeling. After pelleting cellular debris by centrifugation, the protein was affinity purified using TALON cobalt affinity resin (Clontech Laboratories, Mountain View, CA) followed by size exclusion chromatography on Superdex 200 10/30 GL size exclusion column (GE Healthcare, Little Chalfont, UK) into size exclusion buffer (SEB): 150 mM NaCl, 20 mM MOPS pH 7.0, 5 mM analytical-grade DM. Molar extinction coefficients for WT and RCLC are 46,020 M^-1^ cm^-1^ and 49,630 M^-1^ cm^-1^ respectively, and molecular weights for WT and RCLC is 52,000 g/mol and 49,630 g/mol respectively.

### Preparation of lipids

Detergent solubilized lipid micelles were prepared as described before (4). Briefly, a desired amount of *E. coli* polar lipid extract (EPL; Avanti Polar Lipids Inc., Alabaster, AL) in chloroform (25 mg/mL stock) was dispensed in a glass vial. The chloroform was evaporated under a continuous stream of 0.22 μm filtered Ultra High Purity N_2_ gas (Airgas), and then the lipids were washed in pentane and then dried while rotating the vial, approximately 10-12 minutes, leaving a thin film of lipids along the walls and the bottom of the tube. After repeating the washing step twice, the lipid film was resuspended in Dialysis Buffer (DB: 300 mM KCl, 20 mM citrate pH 4.5, adjusted with NaOH) to a final concentration of 20 mg/mL and 35 mM of CHAPS was added. The lipid/detergent mixture was solubilized using a cup-horn sonicator (Qsonica) until the sample achieved a homogenous translucent appearance.

### Protein labeling and reconstitution

After purification by size-exclusion chromatography, 10 μM protein was reacted with 50 μM of either Cy3-, or Cy5-, or a 1:8 mixture of Cy3- and Cy5-(Cy3/5) maleimide in DMSO (Lumiprobe Corporation, Hunt Valley, MD) for 10 minutes, then quenched with 5 mM cysteine from a 100 mM cysteine stock in SEB, pH adjusted to 7.0. The free dye was removed by re-binding the protein to a 0.2 mL cobalt affinity column (Talon), followed by excessive washing with 10-15 CV of cobalt column wash buffer (CoWB: 100 mM NaCl, 20 mM Tris, 5 mM DM, pH 7.5), and then elution with CoWB containing 400 mM imidazole, pH 7.5. As a final step, the imidazole was removed by running the eluted sample on a 3 mL Sephadex G-50 size exclusion column (Sigma-Aldrich). The labeling yield was quantified by placing the sample in a 1 cm quartz cuvette in a Nanodrop 2000c UV-VIS spectrophotometer and measuring the absorbance spectrum between 190-840 nm. The protein concentration in the presence of Cy5 (or Cyanine 5) is calculated as:

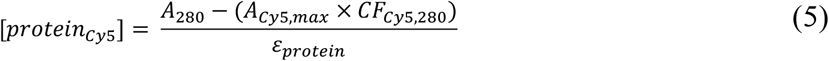

where *A*_280_ is the absorbance at 280 nm, *A*_*cy*5,*max*_ is the peak absorbance of Cy5 ≈ 653 nm, *CF*_*cy*5,280_ = 0.017 (*CF*_*cy*5,280_ = 0.05 for Cyanine 5), is the correction factor for the absorbance of Cy5 at 280 nm, and *ε*_*protein*_ is the extinction coefficient for the protein at 280 nm. The subunit labeling yield, *P*_*Cy5*_, is calculated as:

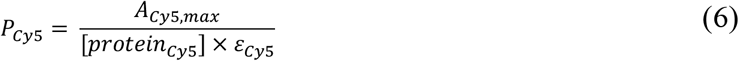

where, *ε*_*Cy5*_ = 2.5 × 10^5^ M^-1^ cm^-1^ is the extinction coefficient for Cy5 at 653 nm. For the Förster Resonance Energy Transfer (FRET) studies, protein was labelled with Cy3-maleimide (Lumiprobe) or simultaneously co-labelled with Cy3- and Cy5-maleimide. For Cy3-labeling alone, the same procedure was followed, except that the quantification is corrected for Cy3 contribution at 280 nm. Thus, the protein concentration in the presence of Cy3 is calculated as:

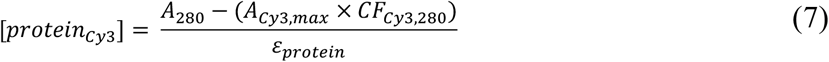

where *A*_*cy*3,*max*_ is the peak absorbance of Cy3 ≈ 555 nm, and *CF*_*cy*3,280_ = 0.08 is the correction factor for the absorbance of Cy3 at 280 nm. The subunit labeling yield, *P*_*Cy3*_, is calculated as:

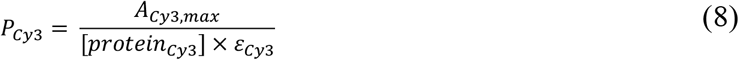

where, *ε*_*Cy*3_ = 1.5 × 10^5^ M^-1^ cm^-1^ is the extinction coefficient for Cy3 at 555 nm. For labeling of Cy3 and Cy5 simultaneously, there are two correction factors to consider, Cy3 and Cy5 absorbance at 280, as well as the contribution of Cy5 absorbance in the Cy3 peak. Thus, the protein concentration in the presence of Cy3 and Cy5 is calculated as:

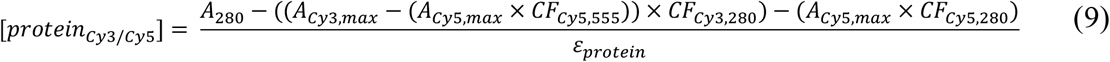

where *CF*_*cy*3,555_ = 0.08 is the correction factor for the absorbance of Cy5 around the Cy3 peak. The subunit labeling yield of Cy3 in the presence of Cy5, *P*_*Cy3:Cy5*_, is calculated as:

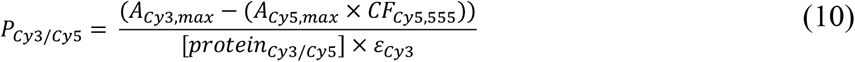

and the subunit labeling yield of Cy5 in the presence of Cy3, *P*_*Cy5:Cy3*_, is calculated as:

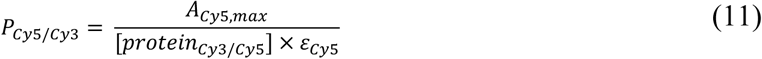

Altogether, we obtained the following Cy5, Cy3, and co-labeled Cy3/5 labeling yields for WT CLC-ec1: *P*_*Cy5*_ = 0.72 ± 0.01 (n = 9), *P*_*Cy3*_ = 0.79 ± 0.01 (n = 9), *P*_*Cy5*_/*P*_*Cy3*_ = 6.0 ± 0.4 (n = 8) (*P*_*Cy3*_ = 0.11 ± 0.01, *P*_*Cy5*_ = 0.63 ± 0.02, respectively). For RCLC CLC-ec1, we obtained the following Cy5, Cy3, and co-labeled Cy3/5 labeling yields: *P*_*Cy5*_ = 0.71 ± 0.01 (n = 6), *P*_*Cy3*_ = 0.73 ± 0.05 (n = 6), *P*_*Cy5*_/*P*_*Cy3*_ = 4.8 ± 0.4 (n = 6) (*P*_*Cy3*_ = 0.13 ± 0.01, *P*_*Cy5*_ = 0.63 ± 0.02, respectively).

For the bulk FRET experiments, co-labeled samples were reconstituted at 1 μg/mg, *χ* = 10^−5^ subunits/lipid in the EPL/CHAPS micelles and then dialyzed against dialysis buffer, at 4 °C in the dark, and maintaining a 1000-fold sample to dialysis buffer ratio with 5 buffer changes over 48-72 hours. The proteoliposomes used in the WT_Cy3_ + WT_Cy5_ fusion experiments were also prepared at 1 μg/mg, *χ* = 10^−5^ subunits/lipid in the EPL/CHAPS micelles, in separate volumes that would correspond to a final acceptor to donor ratio that matches the co-labelled *P*_*Cy5*_*/P*_*Cy3*_ ratio. For example, if *P*_*Cy5*_/*P*_*Cy3*_ = 6 for the co-labeled sample, and *P*_*Cy3*_ = 0.80 and *P*_*Cy5*_ = 0.65 for the independently labeled samples, then 100 μL of WT_Cy3_ and 747 μL of WT_Cy5_ liposomes were prepared and dialyzed separately, and the full volumes of each sample after dialysis were combined for the “fused” samples upon freeze-thaw. This resulted in fused samples with *P*_*Cy5*_/*P*_*Cy3*_ ∼ 6, even if there was variation in the dilution of the lipid concentrations in each cassette during dialysis. For freeze-thawing to form MLVs, proteoliposomes or empty liposomes were retrieved from dialysis, frozen and thawed by alternating between -80 °C and room temperature for 15 and 20 minutes respectively, for a total of two times and then stored at -80 °C until further use. Room temperature is defined as ambient laboratory temperature around 21-23 °C.

For the photobleaching experiments, WT-Cy5 protein was mixed with 20 mg/mL EPL/CHAPS micelles at 0.0001, 0.001, 0.01, and 0.1 μg/mg, corresponding to *χ* =10^−9^, 10^−8^, 10^−7^ and 10^−6^ subunit/lipid mole ratio. In addition, “empty” EPL liposomes were prepared in parallel to use in the freeze-thaw fusion/dilution studies. To avoid any possibility of contamination, samples were dialyzed in separate buckets.

### Bulk FRET measurements in multi-lamellar vesicles

Förster Resonance Energy Transfer (FRET) was measured for fused or co-labeled proteoliposomes in the MLV state using a Fluorolog 3-22 fluorometer with double monochromators on both excitation and emission paths (Horiba Jobin-Yvon, Edison NJ). Samples were excited at 550 nm and spectra was collected from 560 - 740 nm, with a 4 nm slit width, 0.05 s integration time. Typically, 0.3 mL of MLVs were added to a sub-micro quartz cuvette (Starna Cells, Atascadero, CA). Up to four samples were examined in parallel, maintained at a set temperature (22 – 80 °C) in the Peltier controlled 4-position turret holder (Horiba Jobin-Yvon, Edison NJ). The rate of acquisition was set at once every hour for 22, 31, and 37 °C samples, 30 minutes for the 44 °C, once every minute for the 50 and 56 °C, 1.5 minutes for the 62, 68, 74, 80 °C experiments.

The ratiometric FRET signal was calculated as follows:

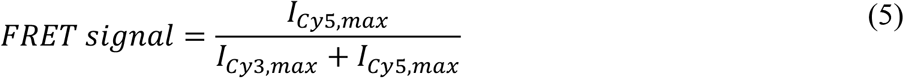

For the FRET reversibility studies, WT_Cy3/Cy5_, WT_Cy3+Cy5_^*fused*^, RCLC_Cy3/Cy5_ or RCLC_Cy3+Cy5_^*fused*^, all at 1 μg/mg, *χ* = 10^−5^ subunits/lipid in EPL, in the MLV state, were retrieved from the cuvette after 48-hours of FRET measurements at 37 °C. The samples were mixed with 5X excess (*v/v*) 2:1 POPE/POPG MLVs containing 1 μg/mg, *χ* = 10^−5^ subunits/lipid of un-labelled WT protein. Typically, 60 μL of labeled sample was mixed with 240 μL of un-labelled sample. The membranes were fused by 3 freeze-thaw cycles, and the FRET scans were resumed on the diluted samples for additional 48 hours at 37 °C. The choice to dilute the samples with 2:1 POPE/POPG proteoliposomes was made to reduce the background signal due to a contaminating fluorescence in EPL.

### Single-molecule subunit capture photobleaching analysis

The details of this method have been described previously (4, 5). Briefly, WT CLC-ec1 is labelled with Cy5 and reconstituted across a wide range of protein densities, from *χ* = 10^−9^ to 10^−6^ subunits/lipid, corresponding to 0.0001 - 0.1 μg/mg of protein per lipid. These proteoliposomes are freeze-thawed (FT) to form large multi-lamellar vesicles (MLVs) where multiple copies of subunits exist within each bilayer even at dilute protein densities. The MLV membranes are incubated at the desired temperature and then the samples are extruded forming small unilamellar vesicles with a defined size distribution. The statistics of subunit capture into the vesicle population follows a Poisson-like distribution and depends on the prior oligomer equilibrium distribution in the larger membranes. We quantify the probability distribution of Cy5-subunit occupancy per liposome by carrying out single-molecule photobleaching analysis on a total internal reflection fluorescence (TIRF) microscope, and then extract the fraction of dimer, *F*_*Dimer*_, of the population by comparing the experimental probability distribution to monomer and dimer control distributions (6). Following this approach, we have been able to show that CLC-ec1 undergoes a reversible dimerization reaction in 2:1 POPE/POPG lipid bilayers, with an equilibrium constant of *K*_*eq*_ = 4.1 × 10^7^ lipids/subunit corresponding to a free energy of *ΔG°*_*2:1 POPE/POPG*_ = -10.9 kcal/mole relative to the 1 subunit/lipid standard state.

In the current studies, reconstituted MLVs were thawed from -80 °C and stored at either room temperature or 37 °C for 72-120 hours in the dark, before extrusion 21-times through a 400 nm nucleopore filter at room temperature. Liposomes were imaged using total internal reflection fluorescence (TIRF) microscopy and single-molecule photobleaching analysis was carried out as described before (4–6). Images were analyzed as described previously using custom image analysis software in MATLAB (30).

In-bilayer dilutions of the samples were prepared by serially diluting WT-Cy5 MLV proteoliposomes into empty EPL MLV liposomes resulting in at 0.0001, 0.001, 0.01, and 0.1 μg/mg, *χ* = 10^−9^, 10^−8^, 10^−7^ and 10^−6^ subunits/lipids respectively, followed by 3 freeze-thaw cycles to fuse the membranes together. The diluted samples were then incubated at the designated temperature (22 – 62 °C) in separate dry baths for indicated period, in the dark, to allow for equilibration of the dimerization reaction. Following this, the MLV samples were cooled down to room temperature and extruded for single molecule photobleaching analysis. For the lower temperatures (4, 11, 18 °C), the 4 °C samples were incubated in the cold room (4 – 5 °C) in the dark, and the samples (11, 18 °C) were incubated in a dry bath in the cold room. The photobleaching experiments were done at 0.001 μg/mg after 2, 4, 6, 16, 20, 45 weeks to check whether the samples reached to dimerization equilibrium after the incubation.

### K_eq_ and van ′t Hoff parameter estimation

The temperature dependent single-molecule photobleaching data was used to estimate *K*_*eq*_ for each sample. First, the mean ± standard deviation from the bootstrapping fits of *K*_*eq*_ were fit with the linear (eq. 3) and non-linear (eq. 6) forms of the van ′t Hoff equations using the linear and non-linear fitting methods in GraphPad Prism. For the linear van ′t Hoff fit, *ΔH°* = -26.0 ± 7.2 kcal mol^-1^ and *ΔS°* = -0.043 ± 0.023 kcal mol^-1^ K^-1^. For the non-linear van ′t Hoff fit, *T*_*0*_ = 300 K is constrained to be constant, *ΔH*_*0*_*°* = 15.5 ± 12.0 kcal mol^-1^, *ΔS*_*0*_*°* = 0.040 ± 0.040 kcal mol^-1^ K^-1^ and *ΔC*_*P*_ = -2.8 ± 0.8 kcal mol^-1^ K^-1^. In addition, we carried out a bootstrapping approach in MATLAB, using a similar script to the bootstrapping estimation of *K*_*eq*_ above, which allowed direct fitting of all four parameters: *T*_*0*_ = 300.0 ± 1.6 K, *ΔH*_*0*_*°* = 9.98 ± 4.06 kcal mol^-1^, *ΔS*_*0*_*°* = 0.077 ± 0.014 kcal mol^-1^ K^-1^ and *ΔC*_*P*_ = -2.47 ± 0.24 kcal mol^-1^ K^-1^ (mean ± standard deviation). The distribution of the bootstrap fitted parameters are reported in **Fig. 4**.

### Functional measurements

Chloride transport assays from 400 nm extruded liposomes were performed as described previously (4, 10). After completion of the bulk FRET measurements, the MLV samples were retrieved from cuvettes and stored in sealed tubes at room temperature, 2-3 days, until the functional assays could be conducted. For 62 ºC experiment, the sample was incubated for 0.1 hrs at 62 ºC in the dry bath to capture the state in equilibrium before the aggregation reaction happens. For higher temperature (68 - 80ºC) samples, the chloride transport assays were incubated for 2.5 ± 0.8 hr at 68 ºC, 1.9 ± 0.2 hr at 74 ºC and 2.7 ± 1.2 hr at 80 ºC so that the plateau is reached in aggregation reaction from bulk FRET measurements.

## Notes

### Competing Interest Statement

The authors have declared no competing interest.

