## Supplementary Figures & Table for "A thermodynamic analysis of CLC transporter dimerization in lipid bilayers"

**Supplementary Table 1.** Temperature dependent $K_D$, $K_{eq}$ and $\Delta G°$ (1 subunit/lipid standard state) values from bootstrap fitting of $F_{Dimer}$ vs. $\chi$* from single molecule photobleaching data. Data represented as mean ± standard deviation from 100 bootstrapped fits. $22^R$ indicates samples where the density was set before reconstitution and incubated at 22 °C.

| Temperature (℃) | $K_D$ (subunits/lipid) | $K_{eq}$ (lipids/subunit) | $\Delta G°$ (kcal/mole) |
|---|---|---|---|
| $22^R$ | $5.8 \pm 5.6 \times 10^{-10}$ | $3.5 \pm 2.6 \times 10^{9}$ | -12.7 ± 0.5 |
| 31 | $5.8 \pm 3.4 \times 10^{-10}$ | $2.5 \pm 1.8 \times 10^{9}$ | -13.0 ± 0.4 |
| 37 | $4.9 \pm 2.4 \times 10^{-10}$ | $2.6 \pm 1.4 \times 10^{9}$ | -13.3 ± 0.3 |
| 44 | $1.1 \pm 0.4 \times 10^{-9}$ | $1.1 \pm 0.6 \times 10^{9}$ | -13.1 ± 0.3 |
| 50 | $2.7 \pm 1.8 \times 10^{-9}$ | $5.2 \pm 3.4 \times 10^{8}$ | -12.8 ± 0.4 |
| 53 | $1.1 \pm 0.3 \times 10^{-9}$ | $9.7 \pm 2.6 \times 10^{8}$ | -13.4 ± 0.2 |
| 56 | $8.5 \pm 3.9 \times 10^{-9}$ | $1.6 \pm 1.2 \times 10^{8}$ | -12.3 ± 0.4 |
| 59 | $5.0 \pm 1.2 \times 10^{-8}$ | $2.1 \pm 0.5 \times 10^{7}$ | -11.1 ± 0.2 |
| 62 | $2.1 \pm 0.7 \times 10^{-7}$ | $5.4 \pm 1.8 \times 10^{6}$ | -10.3 ± 0.2 |

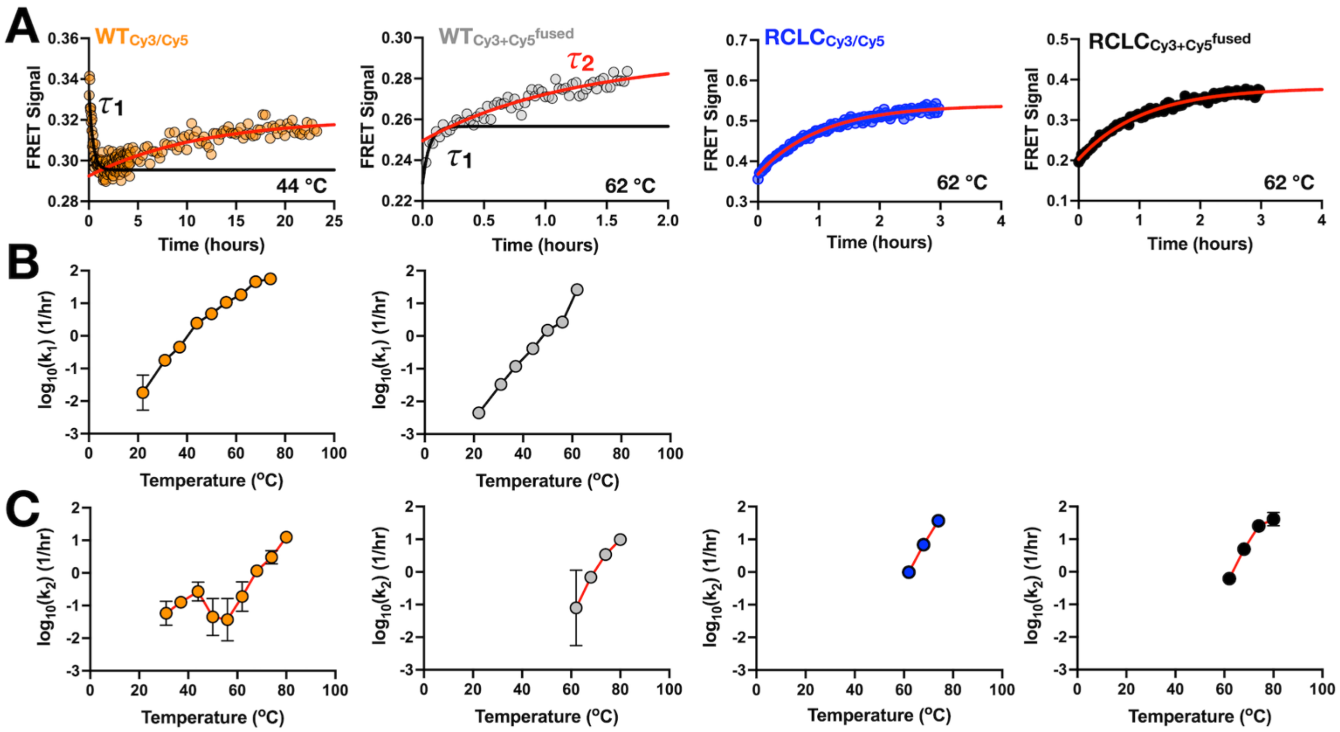


**Figure 2 – Supplement 1. Two-component FRET kinetics in CLC subunit-exchange experiment. (A)** Example raw FRET signals for $WT_{Cy3/Cy5}$ (orange), $WT_{Cy3+Cy5}^{fused}$ (grey), $RCLC_{Cy3/Cy5}$ (blue), and $RCLC_{Cy3+Cy5}^{fused}$ (black). Two components are visible in WT reactions, a faster exponential decay or association indicative of subunit-exchange ($\tau_1 = 1/k_1$, black line) and a slower component corresponding to aggregation that appears at higher temperatures ($\tau_2 = 1/k_2$, red line). The RCLC reactions only contain the $\tau_2$ component. **(B)** Summary of $k_1$ and **(C)** $k_2$ dependencies on temperature (n = 1-5, mean ± sem).

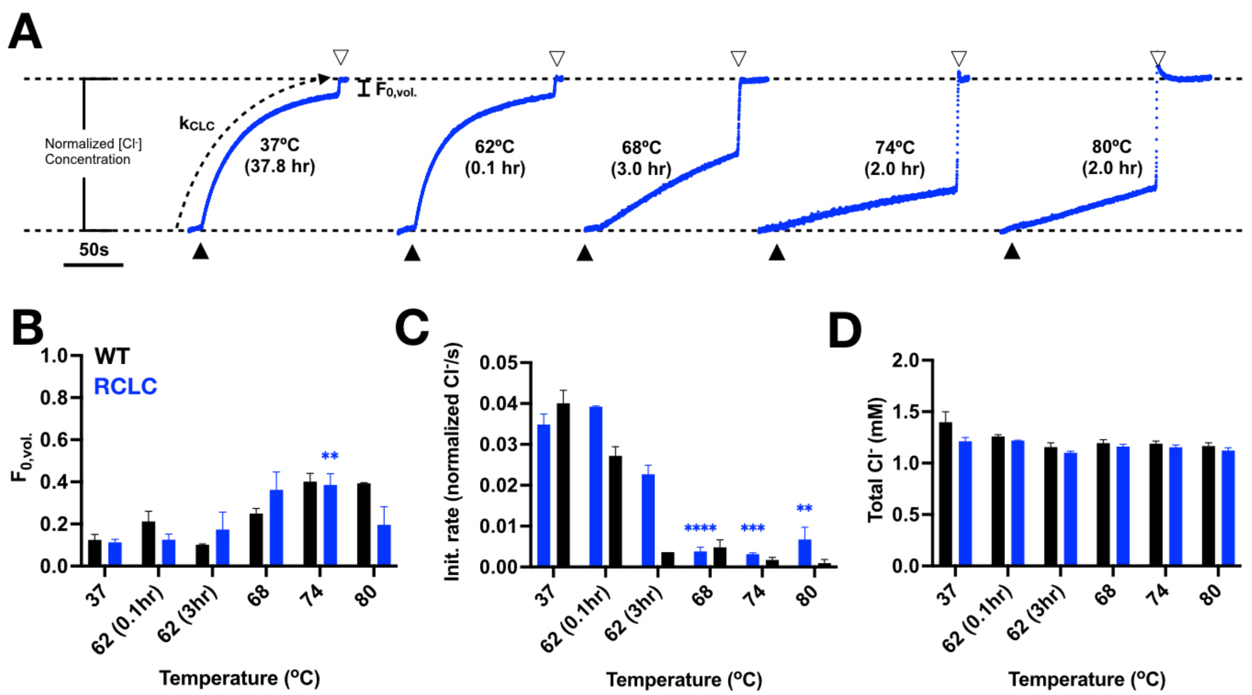


**Figure 3 - Supplement 1. Disulfide cross-linked R230C/L249C CLC-ec1 remains functional for $Cl^-$ transport activity after incubation at elevated temperatures in EPL membranes. (A)** Representative RCLC chloride efflux traces from liposomes reconstituted at $\chi^*$ = 4 x $10^{-6}$ subunits/lipid in EPL from 37 ºC to 80 ºC. The transport activity was measured after incubation 38.7 ± 4.3 hr at 37 ºC, 0.1 ± 0 hr at 62 ºC, 3.0 ± 0 hr at 62 ºC, 3.0 ± 0 hr at 68ºC, 2.0 ± 0 hr at 74ºC, and 2.0 ± 0 hr at 80 ºC ($n$ = 2 - 4). Black triangle indicates the addition of valinomycin and FCCP, white triangle indicates the addition of β-octyl-glucoside. **(B)** Fractional volume of inactive vesicles, $F_{0,vol}$ of RCLC (blue) and WT (black). **(C)** Initial chloride transport rate. **(D)** Total $Cl^-$ concentration in the measurement cell after addition of β-OG. Statistics tests were carried out for RCLC by comparing with 37 ºC data using the unpaired parametric student's t-test.

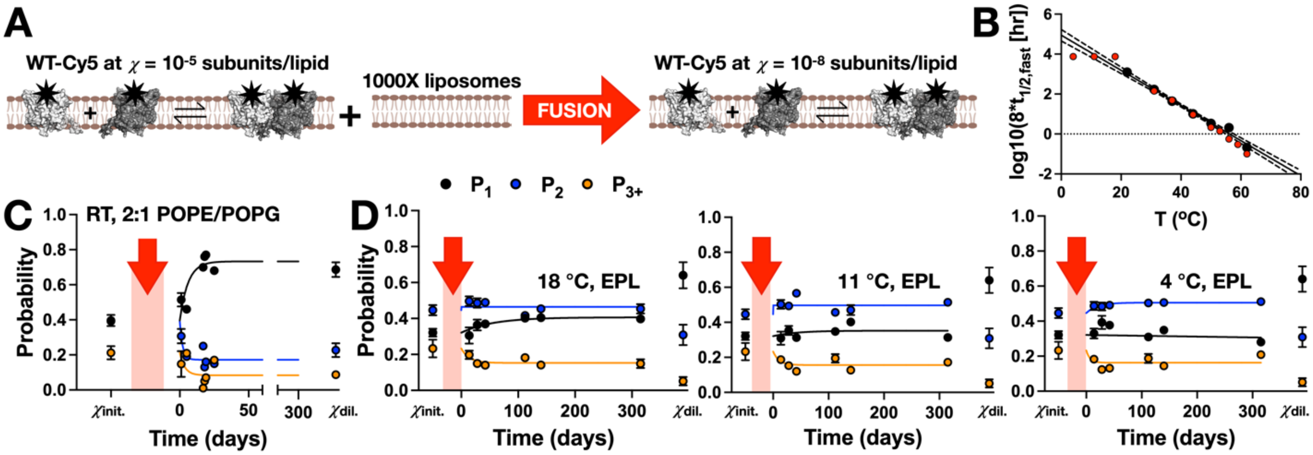


**Figure 4 - Supplement 1. Lack of evidence of CLC-ec1 dimerization equilibration at temperatures below room temperature.** **(A)** Schematic of the in-membrane dilution studies. Dimeric WT-Cy5 reconstituted at $\chi_{rec.}$ = $10^{-5}$ subunits/lipid is diluted 1000-fold, to $\chi$ = $10^{-8}$ subunits/lipid by freeze/thaw fusion with empty vesicles and then incubated at room temperature or lower to monitor the relaxation time-course. **(B)** Predicted incubation times 8 x $t_{1/2}$ from the subunit exchange kinetics, black, and actual incubations times in red. **(C)** Photobleaching step probabilities - $P_1$, $P_2$ & $P_{3+}$ as a function of time for WT-Cy5 in 2:1 POPE/POPG at room temperature, RT ≈ 22 ºC (adapted from Chadda et al., *eLife* 2016), and **(D)** WT-Cy5 in EPL at 18, 11 and 4 ºC. $\chi_{init.}$ corresponds to samples reconstituted at $\chi_{rec.}$ = $10^{-6}$ subunits/lipid and removing the instantaneous dilution effect from multi-occupied vesicles, and $\chi_{dil.}$ corresponds to the expected final dilution reconstituted at $\chi_{rec.}$ =$10^{-8}$ subunits/lipid, both at RT. The red arrow indicates the dilution by freeze/thaw fusion. Data represented as mean ± sem, n = 2-3.
